# From Gradient to Sectioning in CUBE: Workflow for Generating and Imaging Organoid with Localized Differentiation

**DOI:** 10.1101/2022.09.06.506732

**Authors:** Isabel Koh, Masaya Hagiwara

## Abstract

Advancements in organoid culture methods have led to a widespread availability of various *in vitro* mini-organs that mimic native tissues in many ways. Yet, the bottleneck remains to generate complex organoids with body axis patterning, as well as keeping the orientation of organoid samples during post-experiment analysis processes. In this paper, we present a workflow for culturing organoids with gradient using a previously developed CUBE culture device, then sectioning samples with the CUBE to retain information on the gradient direction of the sample. We show that hiPSC spheroids cultured with separate differentiation media on two opposing ends of the CUBE resulted in localized expressions of the respective differentiation markers, in contrast to homogeneous distribution of markers in controls with the same media on both ends. We also describe the processes for cryo and paraffin sectioning of gradient-cultured spheroids in CUBE to retain gradient orientation information. This workflow from gradient culture to sectioning with CUBE can provide researchers with a convenient tool to generate increasingly complex organoids and study their developmental processes *in vitro*.

## Introduction

Pluripotent stem cells (PSCs) and organoids derived from them offer a pragmatic way to model and study the formation of tissues and organs during early human development, as real specimens pose challenging ethical issues (Huch and Koo, 2015; Kim et al., 2020; van den Brink and van Oudenaarden, 2021; Zhu and Huangfu, 2013). Protocols to generate the various organoids that mimic native tissues generally rely on sequentially manipulating the activation or inhibition of signalling pathways such as nodal, hedgehog, notch, Wnt, or BMP at different time points to induce differentiation towards specific lineages, such as in the generation of intestine (McCracken et al., 2011), kidney (Taguchi and Nishinakamura, 2017), and lung (Miller et al., 2019) organoids, epiblasts (Rust et al., 2006; Simunovic et al., 2019), and gastruloids (Veenvliet et al., 2020).

Nevertheless, controlling the growth and differentiation of cells along a body axis remains a challenge in 3D organoid cultures. *In vitro*, anterior-posterior and dorsal-ventral patterning in organoids can arise by cellular self-organisation (Moris et al., 2020; Zheng et al., 2019), or achieved by fusing separately differentiated organoids together as an assembloid such as in cerebral organoids with different brain regions or biliary organoids with liver, biliary tract and pancreas components (Bagley et al., 2017; Koike et al., 2019). The limitation with these methods, though, is that because the cells are cultured in a single uniform medium, the level of control in terms of spatial information supplied to the cells is quite low; all cells within the cluster of cells that make up the organoid receive the same differentiation cues from the signalling molecules in the medium. *In vivo*, on the other hand, concentration gradients of morphogens from other sources also contribute to governing where cells go and what phenotype or pattern they should adopt during development (Christian, 2012; Solnica-Krezel and Sepich, 2012). For example, the formation of the neural tube is reliant on signals from the notochord and non-neural ectoderm layer (Briscoe and Small, 2015), and the nephrons of the kidney develop by exchange of various signals between the ureteric bud and metanephric mesenchyme (Costantini and Kopan, 2010). Thus, there is a need to replicate morphogen gradients in order to generate organoids that more closely resemble the native tissues *in vivo*.

Several engineering technologies have been developed to mimic supplying spatial gradient of signalling molecules to cells *in vitro*. PSCs engineered to express Sonic Hedgehog (Shh) (Cederquist et al., 2019) or agarose beads soaked with morphogens (Ben-Reuven and Reiner, 2020) can be placed close to the developing organoid and the diffusion of molecules from the source to the differentiating cells created a high-to-low concentration gradient of morphogens. Additionally, various modified Transwells and microdevices that enable cells to be cultured with two separate media compartments have also been developed to generate morphogen gradients in opposing directions across the cells (Amadi et al., 2010; Demers et al., 2016; Manfrin et al., 2019; Park et al., 2009; Wang et al., 2017). However, these technologies are not without drawbacks: 1) they require complicated preparation and setup procedures which are not so easy to perform in most biology-based laboratories without specialist skills and equipment, 2) they lack control over the placement and positioning of the sample in the device, and 3) it is difficult to retrieve the sample from the device post-experiment for further analyses without causing a lot of damage to the sample or losing the orientation of the sample. In particular, the ability to retain information on the orientation of the gradient that cells were subject to is critical to ensure proper analysis of the sample.

We therefor aimed to address these issues by developing a simple easy-to-use gradient culture platform to control the differentiation of PSCs into organoids with distinct localized patterns, whilst also retaining sample integrity and gradient orientation for imaging analysis processes. To achieve this, we make use of a CUBE culture device previously developed to enhance the handling ability of samples cultured in ECM hydrogel and repeatability of cell seeding and pattern formation (Hagiwara et al., 2016; Hagiwara et al., 2018). Here, we first show how we can precisely position cells in the desired position in the CUBE. Then, we demonstrate the ease of which gradient can be generated in the CUBE to induce localized differentiation of iPSCs simply by transferring it to a two-compartment gradient chip device. Finally, we present the compatibility of the gradient platform with different post-experiment processing methods (cryo and paraffin sectioning) for imaging and analysis whilst maintaining the gradient orientation information (Fig. 1). With this Gradient-in-CUBE workflow, organoids with controlled body axis differentiation can be achieved, providing ever more complex *in vitro* models in which we can systematically apply and study the effects of morphogen gradients in directing cell differentiation and development into tissues, organs, and eventually body systems to further our understanding of the developmental processes in human.

**Figure 1.**
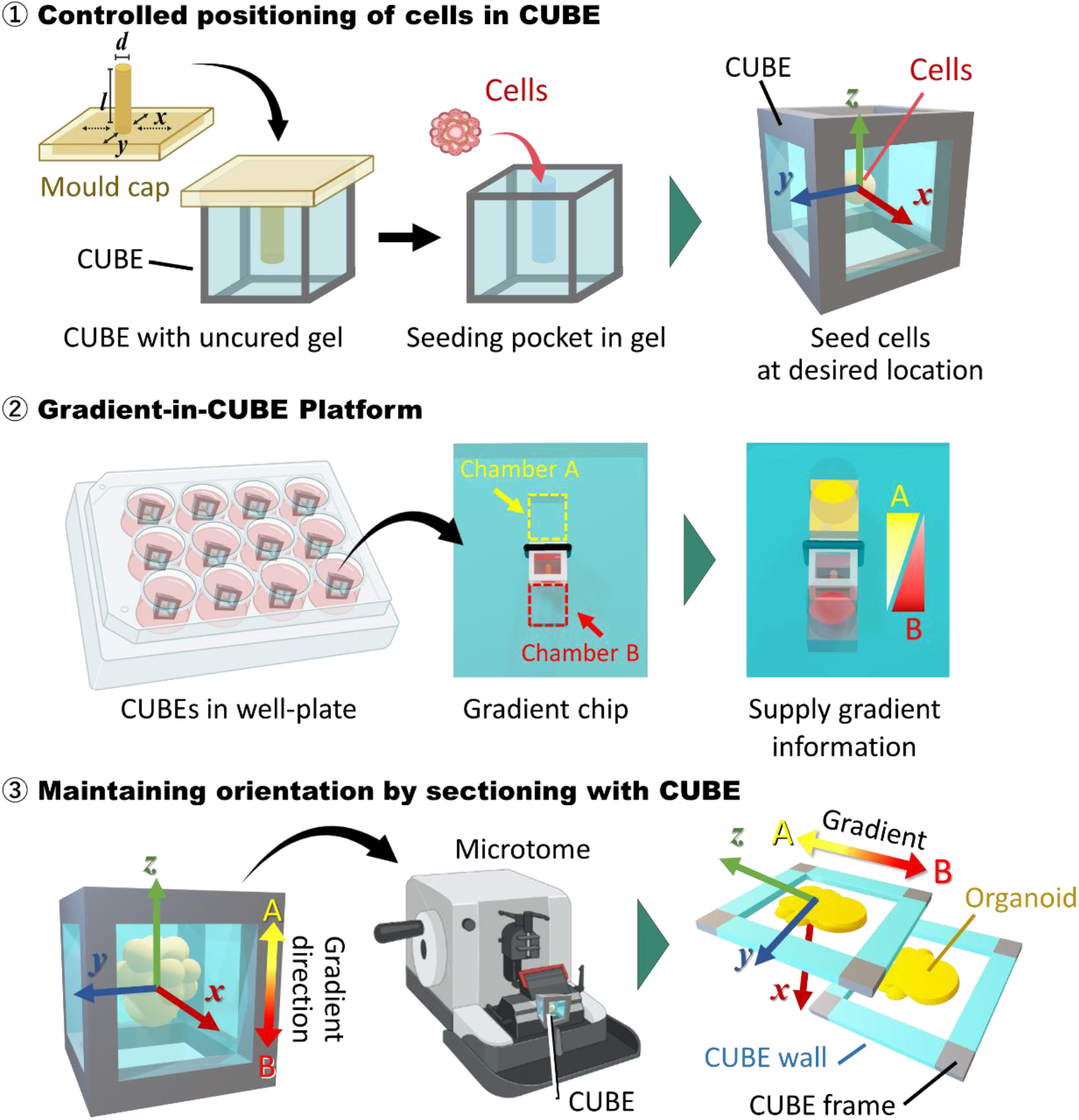
Workflow from gradient culture to sectioning for imaging. The concept for this work was to utilise the CUBE culture device to firstly control the seeding position of cells at the desired location, then transfer the CUBE to a Gradient-in-CUBE chip to culture cells with a morphogen gradient, and finally section the sample with the CUBE to maintain gradient orientation information in the sections.

## Materials and Methods

### hiPSC Maintenance and Spheroid Formation

Human iPSCs (IMR90-4; WiCell Research Institute) were maintained with mTeSR Plus (STEMCELL Technologies, 100-0276) on dishes coated with 9-10 μg/cm^2^ Growth Factor Reduced Matrigel (Corning, 356231), routinely passaged using ReLeSR (STEMCELL Technologies, 05872), and cultured in a 37°C, 5% CO_2_ incubator. Cells were tested for mycoplasma contamination using MycoAlert kit (Lonza, LT07-118). To prepare 96-well plates for spheroid formation, wells were filled with 80 μL/well of 3% agarose (Sigma, A9414) and allowed to solidify. StemFit AK02N (Ajinomoto, RC AK02N) medium plus 10 μM Rock inhibitor (Y-27632; Nacalai Tesque, 08945-84) was then added to the well and incubated. For spheroid formation, 70% confluent cells were dissociated using ReLeSR and resuspended in StemFit +Y27632 medium after centrifugation, then seeded in agarose well-plate. Cells from one 35 mm dish was used to make 5 spheroids. The next day, medium was switched to mTeSR Plus medium without Y27632. Spheroids were transferred to the CUBE device after one day of culture in mTeSR Plus.

### CUBE Fabrication

Two types of acrylic (poly (methyl methacrylate); PMMA) CUBEs were designed for this study using Rhinoceros 3D software. For cryo sectioning, CUBEs were designed with frames of 5 mm lengths and 0.75 mm thickness, with a thicker frame of 1.5 mm on the top side of the cube so that an O-ring can be attached to the cube (Fig. 2Ai). Attachment of the O-ring was necessary to ensure better sealing between the cube and gradient chip to reduce leakage of media around the cube during gradient culture. To make the sidewalls of the Cryo CUBE, polydimethylsiloxane (PDMS; Silpot 184, Dow Toray, 04133124) was first prepared by mixing elastomer base with curing reagent at a 10:1 ratio. Then, a thin layer of the mixture was spread out in a petri dish and degassed to remove air bubbles. The cube frames were placed on the PDMS, then degassed again and baked at 85°C for 30 min to cure the PDMS. Once cured, the PDMS was trimmed from the frames with a scalpel, and the process was repeated to cover the other three sides of the cube, leaving the top and bottom surfaces open (Fig. 2Aii). For paraffin sectioning, CUBEs were designed as a solid cube of 5 mm length with a hollow core of 3.5 × 3.5 mm, with the thickness of the frame being 0.75 mm (Fig. 2Ai). CUBEs for paraffin sectioning had no PDMS sidewalls as PDMS swells in xylene during the paraffinization process. All CUBEs were ordered from machining companies (Cryo CUBEs from Yumoto Electric Inc, Japan, and Paraffin CUBEs from Proto Labs, Japan). Before use, CUBEs were washed twice with ultrasonication, once with MilliQ water and once with isopropanol (IPA), then dried in the oven for 2 hr. For sterilization, the cubes were autoclaved and then dried in the oven at 85°C before use.

**Figure 2.**
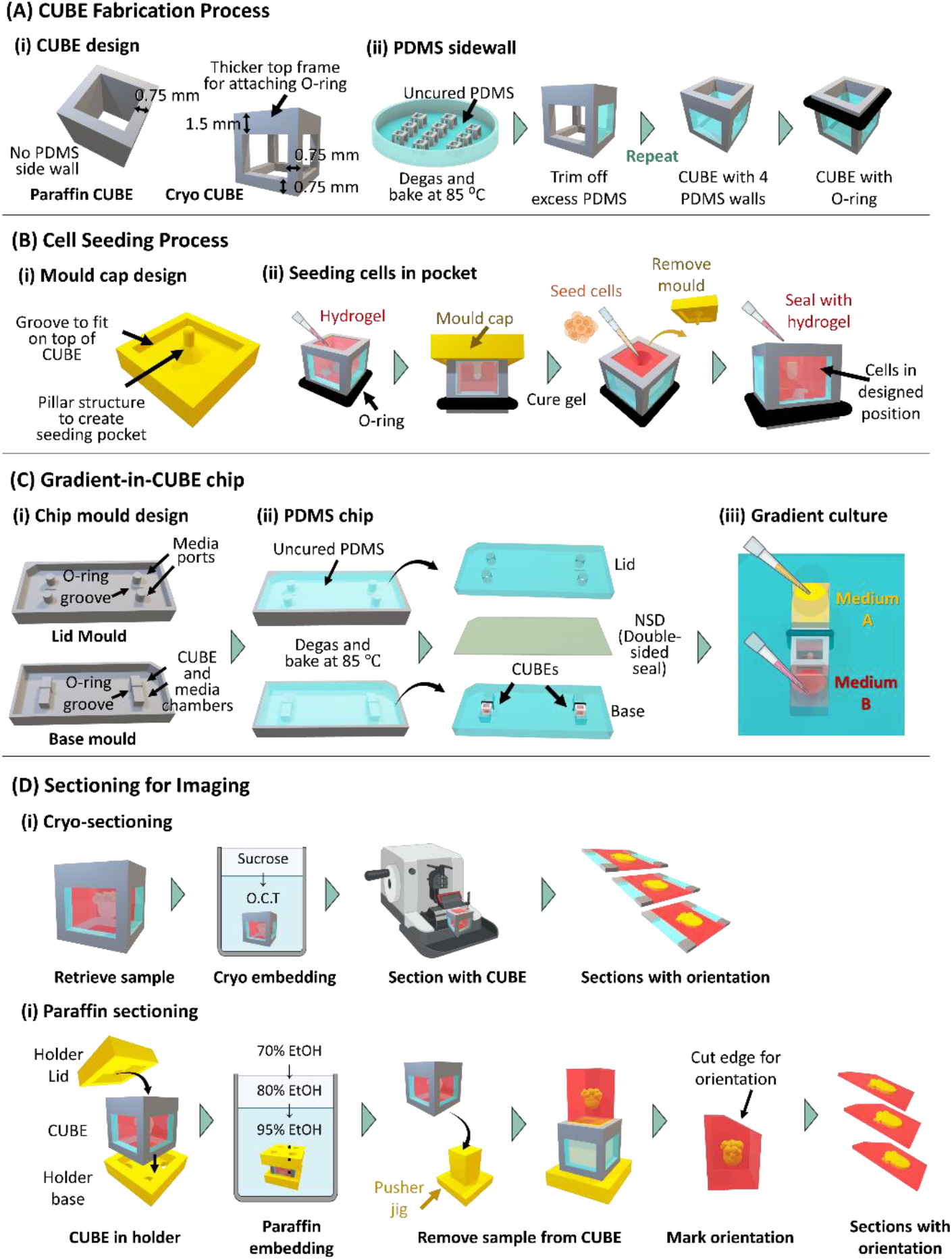
Schematic diagram of fabrication and methods processes. (A) CUBE fabrication process (i) CUBE designs for paraffin and cryo sectioning. (ii) Process to adhere PDMS sidewalls to CUBE device. (B) Cell seeding process (i) Mould cap design. (ii) Process to seed cells in the seeding pocket created by the mould cap in the hydrogel in the CUBE. (C) Gradient-in-CUBE fabrication process (i) Designs for moulds to fabricate the lid and base of the chip. (ii) Process to fabricate PDMS chips from moulds, and adhering the lid to the base by NSD, a double-sided PDMS adhesive seal. (D) Sectioning for imaging. (i) Process to embed samples in cryo medium and sectioning with CUBE. (ii) Process to embed samples in paraffin with CUBE holder, then sectioning with a marked edge to maintain orientation of sample.

### hiPSC Seeding in CUBE

To control the positioning of cells during the initial seeding in the CUBE, a mould cap was designed with a pillar structure at the desired seeding position, and with grooves to ensure the mould fits on top of the cube so that the pillar position can be aligned properly in the cube (Fig. 2Bi). The mould caps were printed using a 3D printer (Agilista 3200, Keyence), and cleaned by ultrasonication twice with IPA after removing excess printing support material, then dried in an oven at 65°C. Before use, the mould cap was dipped in 2-methacrylooyloxtethyl phosphorylcholine (MPC; Lipidure, NOF Corporation, CM5206E) diluted to 5% in isopropanol, then allowed to dry for 1 hr at room temperature (RT). MPC coating helps to ensure smoother detachment of the mould from hydrogel. A nitrile O-ring (AS ONE, 62-3049-63) was attached to the thick part of the acrylic frame for the Cryo CUBE, or approximately 1 mm near one end of the Paraffin CUBE. The CUBEs were placed O-ring side down in a dish, and Matrigel added into the CUBE before placing the mould cap on top of the CUBE and curing the gel in the incubator for 25 min. Once the gel has cured, the mould cap was removed, and hiPSC spheroids were seeded in the pocket created by the mould. Then, additional Matrigel was added to the top of the CUBE and allowed to cure for another 25 min to seal the pocket (Fig. 2Bii). After curing, the CUBE was transferred to a 48-well plate with mTeSR Plus medium and incubated for 2 hrs prior to starting gradient culture.

### hiPSC Differentiation with Gradient-in-Chip

Moulds for making lid of PDMS gradient chips were designed with grooves to fit the O-ring and ports for adding media, while the base was designed to fit the cube, O-ring, and two separate media chambers (Fig. 2Ci). PDMS moulds were ordered from Proto Labs. To make the PDMS chips, uncured PDMS was poured into the moulds, then degassed and baked at 85°C for 1 h to cure the PDMS. Once cured, the chips were removed from the moulds, then washed and sterilized the same way as the CUBEs. To assemble the gradient chip, CUBEs were placed in the chip base with the O-ring fitted into the O-ring groove, then a PDMS double-sided adhesive film (NSD-100, NIPPA) was used to seal the lid and base together (Fig. 2Cii). After assembly, the media chambers on each side of the CUBE were filled with the respective differentiation media (Fig. 2Ciii).

Neuroectoderm differentiation medium comprised 1:1 mixture of KnockOut DMEM/F12 (Gibco, 12660-012) and Neurobasal medium (Gibco, 21103-049) supplemented with 10% KnockOut serum replacement (Gibco, 10828010), 1% MEM non-essential amino acid (Nacalai Tesque, 06344-56), 1% GlutaMAX (Gibco, 35050-061), 1 μM LDN1913189 (Sigma, SML0559), 2 μM SB431542 (Nacalai Tesque, 18176-54), 3 μM CHIR99021 (Nacalai Tesque, 18764-44), 0.1 mM 2-mercaptoethanol (Nacalai Tesque,, 21438-82), and 0.5 μM ascorbic acid (Nacalai Tesque, 03420-52). Mesendoderm basal medium comprised RPMI medium 1640 (Gibco, 11875-093), 1% GlutaMAX, and 1% penicillin-streptomycin (Gibco, 15140122). For mesoderm differentiation, 5 μM CHIR99021 was added to mesendoderm basal medium on Days 0 and 1. From Days 2 to 4, CHIR99021 was withdrawn and replaced with 100 ng/mL bFGF (Nacalai Tesque, 19155-36) and 1 μM all-trans retinoic acid (Stemgent, 04-0021). For endoderm differentiation, 5 μM CHIR99021 was added to mesendoderm basal medium on Day 0, and from Days 2 to 5 CHIR99021 was withdrawn and replaced with 100 ng/mL Activin A (R&D Systems, 338-AC-050/CF). Media change was performed every day by discarding spent media, washing once with DPBS, then replacing with fresh media.

### Fixation and Sectioning for Imaging

At the appropriate timepoint for analysis, the samples were removed from the gradient chip and transferred to a 48-well plate for fixation. Samples were washed twice with DPBS for 5 min each time, fixed with 4% paraformaldehyde for 20 min, then washed twice with DPBS for 10 min each time. For cryo sectioning, samples were soaked in 10%, 15%, and 20% sucrose (Wako, 194-00011) in succession for 1 hr each, then in cryosection embedding medium (Tissue-Tek O.C.T compound; Sakura Finetek, 4583) for 10 min before embedding in embedding medium and freezing at -80°C for 30 min. After freezing, samples were removed from the mould and sectioned as is with the CUBE (Figure 2Di). Sections collected on glass slides were dried with cool air for 30 min, then in an oven at 40°C for another 30 min, followed by washing in MilliQ water to remove excess embedding medium. Then, excess water was removed, and the sections dried at RT for 10 min.

For paraffin sectioning, samples were dehydrated and embedded in paraffin with the following procedure: 70% ethanol for 2 hr, 70% ethanol overnight, 80% ethanol for 1 hr, 95% ethanol for 1 hr, 99% ethanol for 1 hr, 100% isopropanol for 1 hr twice, 100% xylene for 1 hr three times, 1:1 mixture of xylene and paraffin (Nacalai Tesque, 26029-05) overnight at 40°C, and paraffin for 1.5 hr three times at 65°C. To prevent the loss of sample due to shrinkage of Matrigel during the dehydration process, a CUBE holder with lid and base held together by wire was designed to contain the sample in the CUBE (Figure 2Dii). After the final paraffin step, the sample was removed from the CUBE holder and embedded in fresh paraffin. The CUBE was then cut out from the paraffin, and the sample was removed from the CUBE by cutting the paraffin along the inner frame of the CUBE using a scalpel and pressing the CUBE onto a pusher jig (Figure 2Diii). One edge of the paraffinized sample was cut to mark the orientation of gradient direction, then the sample can be sectioned into slices. Deparaffination was performed as follows: 100% xylene for 10 min twice, 99% ethanol for 5 min twice, 95% ethanol for 5 min, 80% ethanol for 5 min, 70% ethanol for 5 min, MilliQ water for 5 min. Heat-induced antigen retrieval was performed by placing the samples in 10 mM citrate buffer (1.8 mM citric acid, 8.2 mM trisodium citrate, pH 6) and heating the buffer until it boils, then cooling it down for 1 min. The boiling and cooling process was repeated 6 times with 10s of boiling followed by 1 min cool down. After the last cycle, samples were cooled for 30 min, then washed with MilliQ water for 5 min, followed by DPBS for 5 min three times.

### Immunofluorescence Staining

Cryo-sectioned samples were permeabilized with 0.5% Triton X-100 for 10 min, followed by three washes with 100 mM Glycine for 10 min each time. Immunofluorescence buffer (IF buffer) was made up of 0.5% Tween20, 2% Triton X-100, and 10% bovine serum albumin (BSA; Sigma, 126615) in DPBS. Blocking was performed by incubating samples with IF buffer with 10% goat serum (Gibco, 16210064) (IF+G) for 30 min, then IF+G with 1% goat anti-mouse IgG (Bethyl Laboratories, A90-116A) for 20 min. Antibodies were diluted according to Table 1, and incubation times were 90 min for primary antibodies and 50 min for secondary antibodies. After each antibody incubation, samples were washed with IF buffer for 15 min three times. Nuclei were stained with DAPI for 20 min, then washed with DPBS for 5 min three times.

**Table 1.** List of antibodies for immunofluorescence staining.

| Reagents | Source and Identifier | Dilution |
| --- | --- | --- |
| Anti-Sox2 | Abcam, ab79351 | 1:200 in IF+G |
| Anti-Nestin | Abcam, ab105389 | 1:200 in IF+G |
| Anti-Brachyury | R&D Systems, AF2085 | 1 μg/mL in IF+0.1% BSA |
| Anti-FoxA2 | R&D Systems, AF2400 | 1 μg/mL in IF+0.1% BSA |
| Anti-Sox17 | R&D Systems, AF1924 | 1 μg/mL in IF+0.1% BSA |
| Donkey anti-goat AF555 | Invitrogen, A32816 | 1:200 in IF+0.1% BSA |
| Goat anti-mouse AF488 | Invitrogen, A32723 | 1:200 in IF+G |
| Goat anti-rabbit AF488 | Invitrogen, A32731 | 1:200 in IF+G |
| DAPI | Invitrogen, D1306 | 600 nM in DPBS |

### FITC-Dextran

To track the formation of gradient over a period of time, CUBE filled with 1.5% agarose was placed in the gradient chip with 10 μM 40kDa fluorescein isothiocyanate (FITC)-dextran (Sigma, FD40S) in DPBS at one end of the CUBE, and DPBS only on the other end. Time-lapse imaging was performed at 10 min interval using a fluorescence microscope (BZX-700, Keyence). After 24 hr, both FITC-dextran and DPBS were discarded, and the media chambers were washed once with DPBS before replacing with fresh FITC-dextran and DPBS. Then, time-lapse imaging was continued for another 24 hr. FITC intensity was measured from time-lapse images using ImageJ software.

## Results and Discussion

Gradients of morphogen signalling contribute to cell fate specification and patterning in developing tissues (Rogers and Schier, 2011), and recapitulating this phenomenon *in vitro* could promote the development of increasingly complex organoids models, which is difficult to achieve in organoids that are cultured in a single uniform medium in a well-plate and rely mainly on self-organisation for patterning (Hofer and Lutolf, 2021).

Although there have been many reports on methods to culture organoids with morphogen gradients, the complicated device setup and poor tissue handling ability, as well as maintenance of gradient orientation during analysis, limit their widespread adoption by the organoid community. By taking advantage of the easy handling ability of the CUBE device, we established a workflow from culturing organoids with gradient to imaging samples with gradient orientation, with the following processes: 1) control the initial seeding position of cells in the CUBE so that cells receive consistent gradient information in each experiment, 2) generate a gradient of morphogen signalling along an axis of the cell sample, and 3) section the sample with the CUBE to retain information of the gradient direction.

### Design of CUBE

Material choice for the CUBE had to be carefully considered to accommodate the requirements of each procedure, particularly as the paraffinization process entails the use of organic reagents. The CUBE frame was switched from the previously used polycarbonate to acrylic material, which is more resistant to xylene treatment for short periods of time compared to polycarbonate. To prevent the leakage of media around the CUBE when it is placed in the gradient chip, the attachment of an O-ring to the CUBE was necessary to seal the gaps that form between the hard material of the CUBE and the softer PDMS chip. Hence, CUBE frame was designed with a thicker frame (1.5 mm) on one side to accommodate the O-ring (Fig. 2Ai). PDMS, a clear silicone-based biocompatible material was used as the sidewall material to restrict movement of morphogen-containing media through the CUBE to only the top and bottom sides of the CUBE, thereby generating a gradient in only one axis of the CUBE. The process to adhere PDMS walls to the CUBE is illustrated in Fig. 2Aii and described in the methods section. However, because PDMS swells when exposed to xylene, CUBEs for paraffin sectioning were designed without sidewalls (Fig. 2Ai). Although this slightly reduces the clarity of samples, a fully acrylic frame still afforded sufficient visibility under brightfield.

### Precise Seeding Positioning in CUBE

The precise placement of cells in the CUBE is important to ensure consistency in the gradient signalling that cells receive. For example, the gradient information sensed by cells that were placed too close to one end of the CUBE will be different from that sensed by cells in the centre of the CUBE. To control the seeding position of cells in the CUBE, a mould cap with a pillar structure to create a seeding pocket at the desired position in the hydrogel in the CUBE, as well as grooves to ensure the precise fitting of the mould cap on the CUBE was designed (Fig. 2Bi). The process to make a seeding pocket and seed cells in the pocket is illustrated in Fig. 2Bii and described in the methods section. By utilizing this seeding method, cells can always be seeded in the desired location with low variation, compared to manually positioning the cells without a guide which results in inaccurate seeding with high variation (Fig. S1).

### Gradient Generation and Localized Differentiation Using Gradient-in-CUBE Chip

To establish a gradient across the length of the CUBE, a Gradient-in-CUBE chip, comprising a base component and a lid component, was designed and fabricated by a moulding process. The mould for the base contained a compartment for the CUBE and two separate media chambers, as well as a groove to fit the O-ring; the mould for the lid has two ports for adding media to the separate media chambers (Fig. 2Ci). The process to make the chip is illustrated in Fig. 2Cii-iii and described in the methods section.

Time-lapse imaging was performed to monitor the progress of an FITC-dextran gradient formation in the CUBE (Fig. 3A), and the intensity of FITC (*I*) across the centre region (*S*) of the CUBE (*x – x’*) was measured (Fig. 3B). The gradient was calculated as the ratio of *I*_*x*_/*I*_*x’*_, where a value of one shows no gradient and a higher ratio represents a steeper gradient (Fig. 3C). A gradient begins to form as FITC-dextran molecules move from the side where it is highly concentrated, to the opposite side where there is no FITC-dextran. The steepness of the gradient increased over the first 4 hours as the gradient forms, but as FITC-dextran accumulates in the opposite DPBS side and in the gel in the CUBE, the steepness of the gradient gradually decreases. Nevertheless, after approximately 24 hours, the gradient stabilizes and can be sustained without falling to equilibrium. Given that a gradient across a scale of a single or a few cells’ length is sufficient to provide positional information to cells (Matos et al., 2020; Vetter and Iber, 2022), we postulated that the gradient generated in the Gradient-in-CUBE chip should be sufficient to induce localized differentiation on opposite ends of a spheroid. The interplay between morphogen and its inhibitor is also critical for patterning in tissues. For example, the Wnt and Nodal antagonists Dkk1, Lefty-1, and Cer-1 in the anterior parts of the embryo restricts Wnt/Nodal to the posterior end of the embryo to establish the anterior-posterior axis (Carlson, 2018).

**Figure 3.**
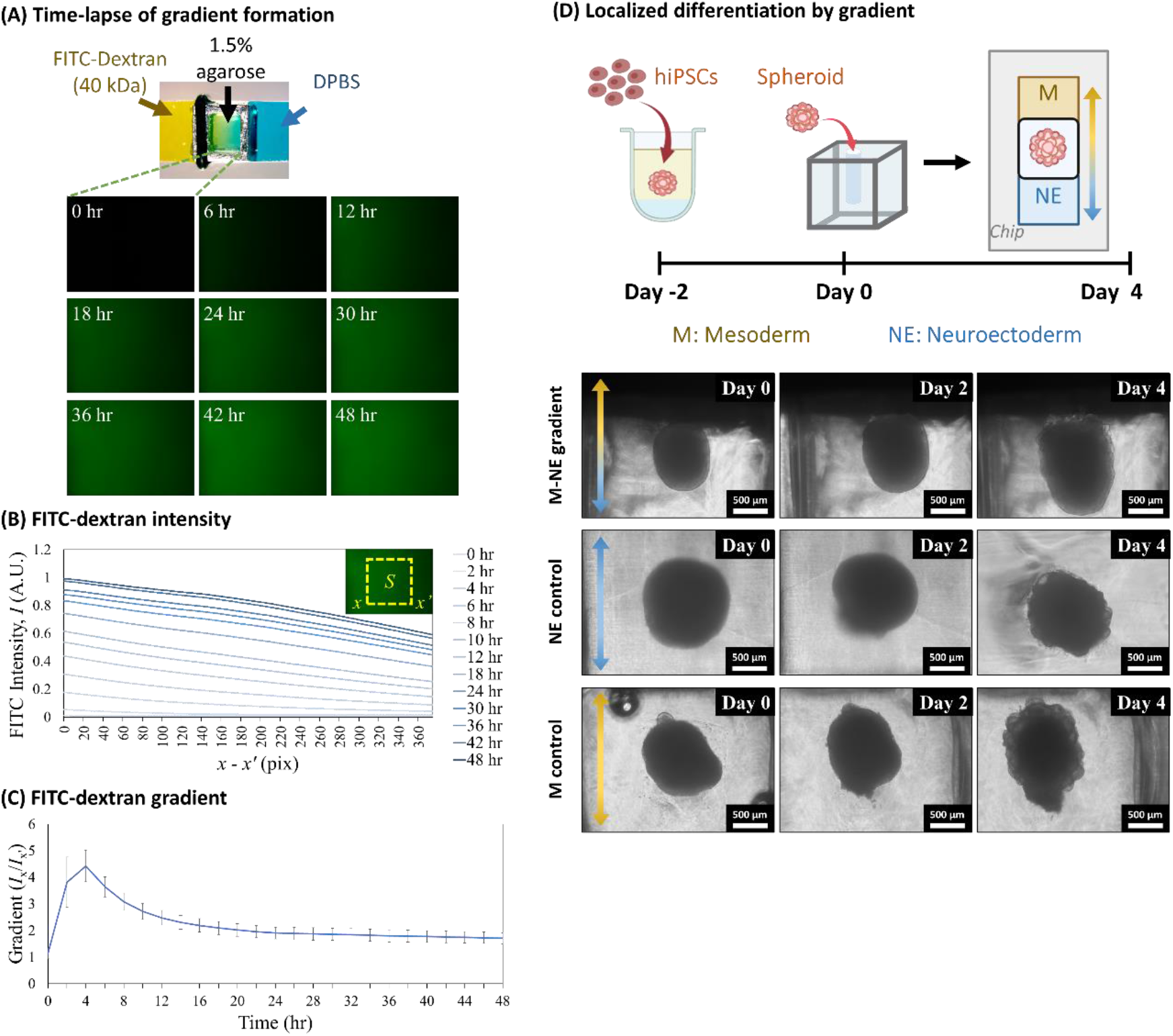
Generating gradient and localized differentiation using the Gradient-in-CUBE Chip. (A) Time-lapse imaging of FITC-dextran gradient forming over a 48 hr period. (B) Intensity of FITC-dextran (*I*) measured over the central region *S* of the time-lapse images from *x* to *x*’ shows the formation and maintenance of a gradient in the CUBE. (C) Gradient calculated as the ratio of *I*_*x*_/*I*_*x’*_ shows gradient increased and peaked at 4 hr, then gradually decreases and remains stable from approximately 24 hr. (D) Differentiation of hiPSC spheroid with mesoderm-inducing medium (M) on one side and neuroectoderm-inducing medium (NE) on the opposite side results in a spheroid with different morphology on either end of the spheroid, whereas the morphology is relatively uniform in M or NE only controls.

To demonstrate the application of the Gradient-in-CUBE chip to differentiate a single spheroid into two localized regions, we supplied a neuroectoderm differentiation medium (NE) to one end of the CUBE, and mesoderm differentiation medium (M) on the opposite side. The mesoderm medium (based on (Lam et al., 2014)) contained high concentrations of Wnt activator CHIR99021, whereas the neuroectoderm medium (based on (Bianchi et al., 2018)) contained lower Wnt activator along with Nodal/Activin inhibitor SB431542 to counteract mesoderm differentiation on the neuroectoderm side. After 4 days of differentiation, morphological differences could be observed at both ends of the spheroid, with the mesoderm side having a bumpier and more protruding feature, compared to smoother feature of the neuroectoderm side. In contrast, control spheroids showed rather uniform morphological features (Fig. 3D).

### Maintaining Orientation during Sectioning and Imaging

It is often difficult to visualize the expression of cell pluripotency or differentiation markers in larger scale 3D samples like organoids, due to the limited range of laser penetration, as well as the low sensitivity of low magnification objective lenses and short focal lengths of high-sensitivity lenses. Hence, organoids are often embedded cryo medium or paraffin and sliced into thin sections for staining and imaging. However, the orientation of the sample is often lost after retrieving the sample from prevailing gradient-generating devices. The advantages of the CUBE device are not only that the cells are contained within the CUBE and can be retrieved without causing damage to the sample, but also that the orientation of the gradient can easily be marked to be recognized later. Here, we show that the samples in the CUBE can be processed for cryo and paraffin sectioning.

For cryo sectioning, the thicker frame on one end of the CUBE where the O-ring was attached acts as an orientation marker. After retrieving the sample from the Gradient-in-CUBE chip, the CUBE can simply be soaked in sucrose and cryo embedding medium before freezing. Once frozen, the CUBE is sliced together with the sample, with the CUBE frame and PDMS outer wall acting as a reference to preserve sample orientation (Fig. 2Di, Fig. 4A). From immunofluorescence staining of hiPSC spheroid differentiated with NE-M gradient, localisation of neuroectoderm marker Sox2 and mesoderm marker Brachyury were observed on the respective ends of the spheroid, whereas in the control samples, both markers were expressed uniformly throughout the spheroid (Fig. 4B). Thus, we demonstrated a method to preserve the orientation information of spheroid differentiated with gradient by sectioning the sample as is in the CUBE. A downside to this method, however, is that the microtome blade gets damaged by repeated cutting of the hard acrylic material and may need to be replaced frequently.

**Figure 4.**
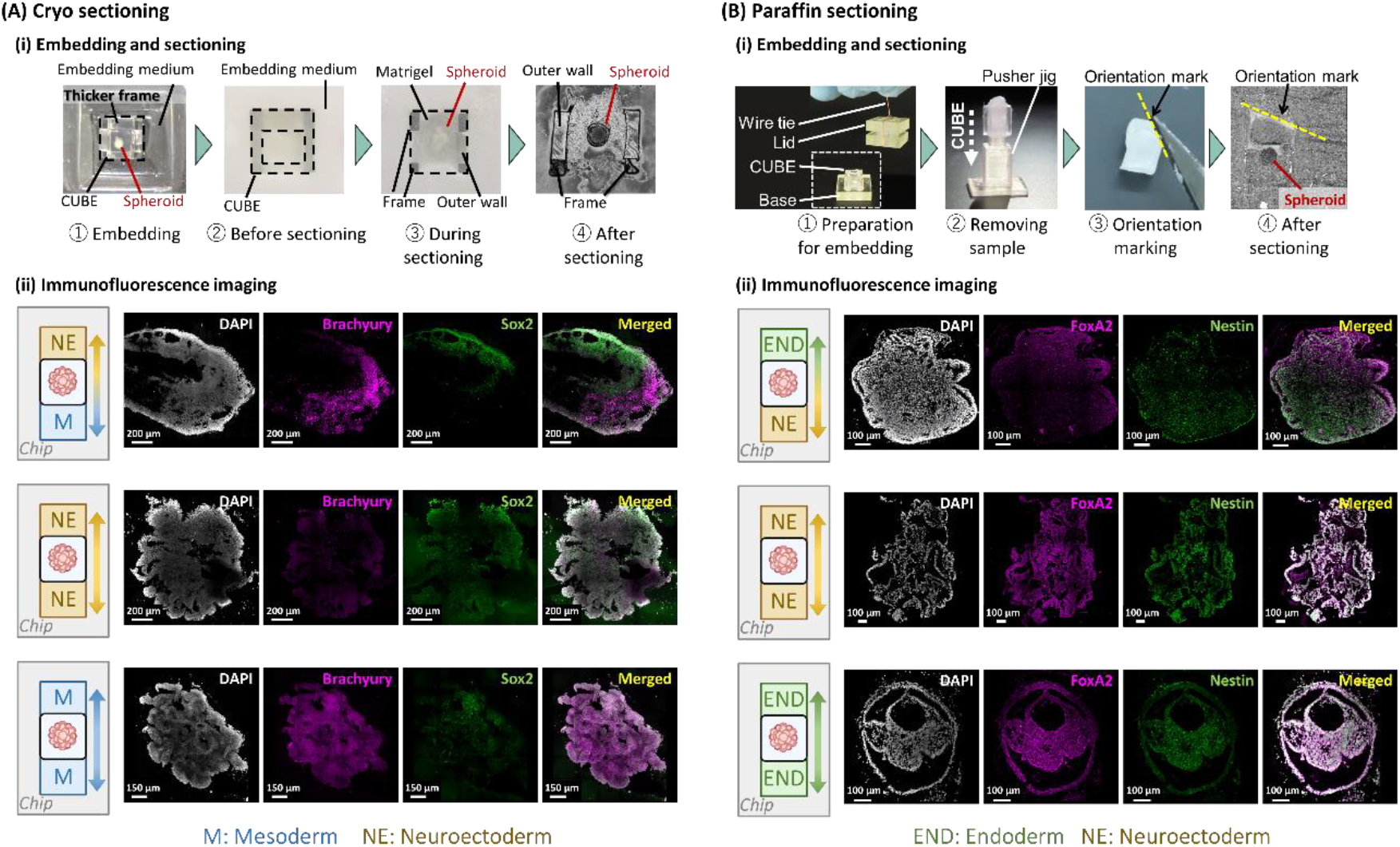
Immunofluorescence imaging after cryo and paraffin sectioning. (A) Cryo sectioning and imaging. (i) Steps to embed and section frozen samples with CUBE, making use of the CUBE frame as a reference marker for sample orientation. (ii) Immunostaining of hiPSC spheroid differentiated with M-NE gradient showed localized expression pattern of mesoderm (Brachyury) and neuroectoderm (Sox2) markers, whereas NE only and M only controls showed uniform distribution of the markers. (B) Paraffin sectioning and imaging. (i) Steps to embed and section paraffin samples with CUBE holder to prevent sample loss, then removing sample from CUBE and cutting an edge of the sample to mark the orientation. (ii) Immunostaining of hiPSC spheroid differentiated with endoderm (END)-NE gradient showed localized expression pattern of endoderm (FoxA2) and neuroectoderm (Nestin) markers, whereas NE only and END only controls showed uniform distribution of the markers.

For paraffin sectioning, some extra steps have to be taken to preserve the integrity and gradient orientation of the sample in the CUBE (Fig. 2Dii, Fig. 4Bi, Fig. S2). During the initial dehydration process prior to paraffin embedding, the hydrogel loses a lot of volume and shrinks in the CUBE. To avoid the risk of losing the sample that may detach from the CUBE, a CUBE holder comprising a lid and a base was designed to cover most of the top and bottom open surfaces of the CUBE but still allow reagents to pass through in and out of the sample. The CUBE in the holder was then tied with a wire to keep them together. After the paraffinization process, the sample was removed from the CUBE using a pusher jig, and the orientation marked by cutting an edge on one corner of the sample. To demonstrate the application of the paraffin method, hiPSC spheroid was differentiated with NE and endoderm (END) differentiation (based on (Lam et al., 2014)) media on either side of the CUBE. Immunofluorescence staining of neuroectoderm markers Nestin and Sox2, and endoderm markers FoxA2 and Sox17, showed localization on the NE and END sides of the spheroids, respectively (Fig. 4Bii and Fig. S3). On the other hand, control samples without gradient culture showed uniform expression of the markers throughout the spheroid.

## Summary

Culturing 3D organoids with gradient has long been a challenge both in terms of complicated gradient device setup and maintaining the orientation of the gradient during analysis processes. In this paper, we present a gradient culture-to-imaging workflow that utilizes the modular CUBE culture device to easily set up a Gradient-in-CUBE device that requires no complicated microfluidic setup. Furthermore, post-experiment, the sample can be easily removed from the device without damaging the sample whilst retaining information on the direction in which the gradient was formed. An interesting point to consider in future works, however, is the potential contribution of differences in mechanical stimulation that cells experience depending on the size of the spheroids in relation to the size of the seeding pocket created by the mould cap, as the sealing of the pocket with additional hydrogel may introduce some variation in gel stiffness due to excess media in the pocket when spheroids are seeded in it. The Gradient-in-Cube platform also has the potential to be adapted to generate gradients in two axes, for example, the anterior-posterior and dorsal-ventral axes in the same organoid, which is work that is currently being undertaken. Hence, the workflow from generation of organoid with localized differentiation to analysis with gradient information could provide researchers a user-friendly method to develop increasingly complex organoids to expand our understanding of various developmental processes.

## Acknowledgements

This research was supported by funds from JSPS KAKENHI Grant numbers JP21H01299 and JP21K18048. Some illustrations were generated using Biorender application.

## Data Availability

All data available upon reasonable request.

## Figures

**Supplementary Figure 1.**
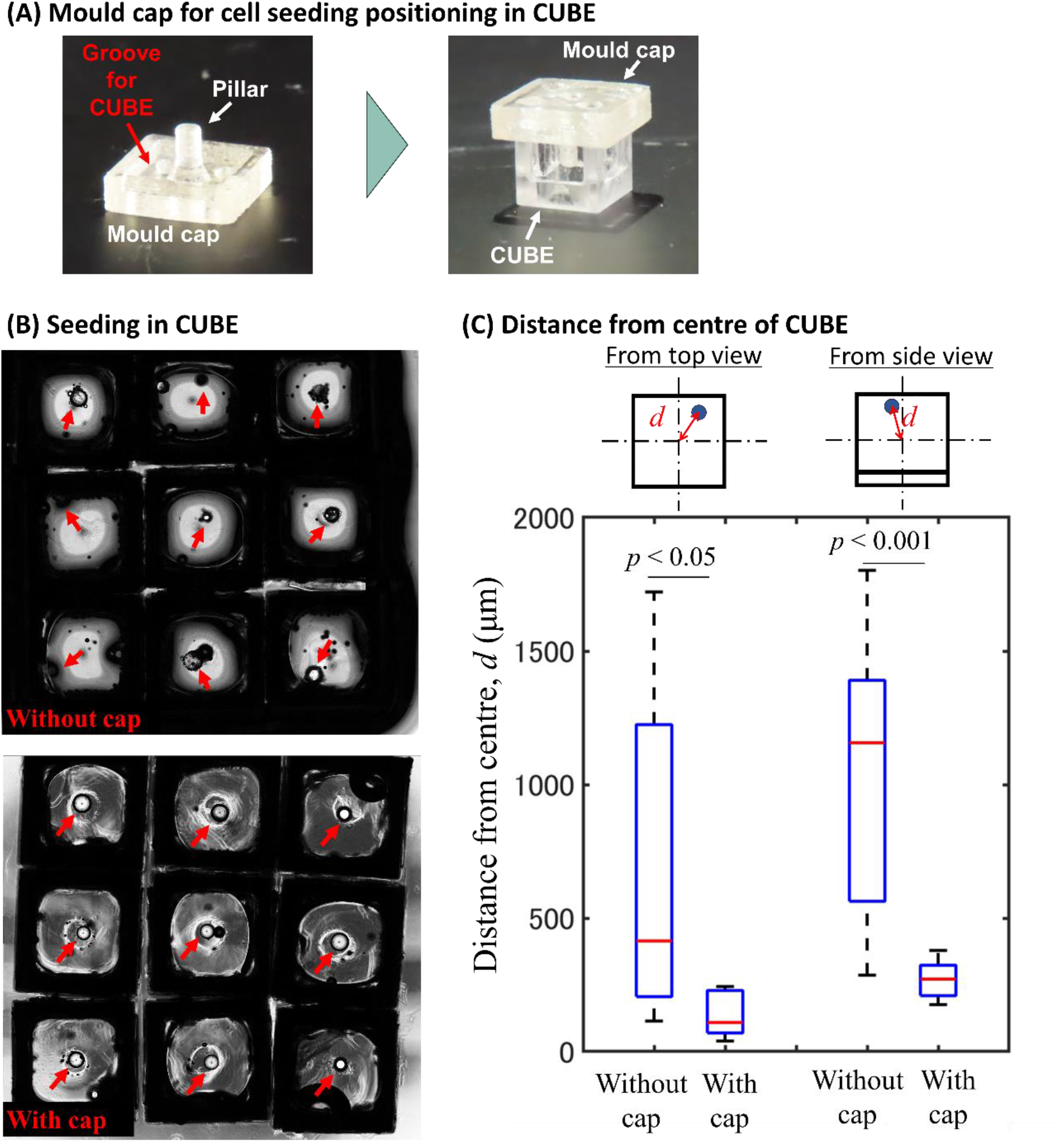
Controlling the seeding position of cells in the CUBE. (A) By placing a mould cap in the hydrogel in the CUBE, a seeding pocket is formed in the gel after it has cured and the mould cap is removed. (B) Top view of brightfield images of PDMS spheres seeded in CUBEs with and without the use of the mould cap to represent hiPSC spheroid seeding. Red arrows indicate the position of the PDMS spheres. (C) Measurements of the distance of PDMS sphere from the centre of the CUBE, from both the top view and side view of the CUBE, show that seeding in the pocket created by the mould cap resulted in consistent seeding position close to the centre of the CUBE compared to manual seeding without a guide. *p* value calculated by one-tailed equal variance t-test; *n* = 8.

**Supplementary Figure 2.**
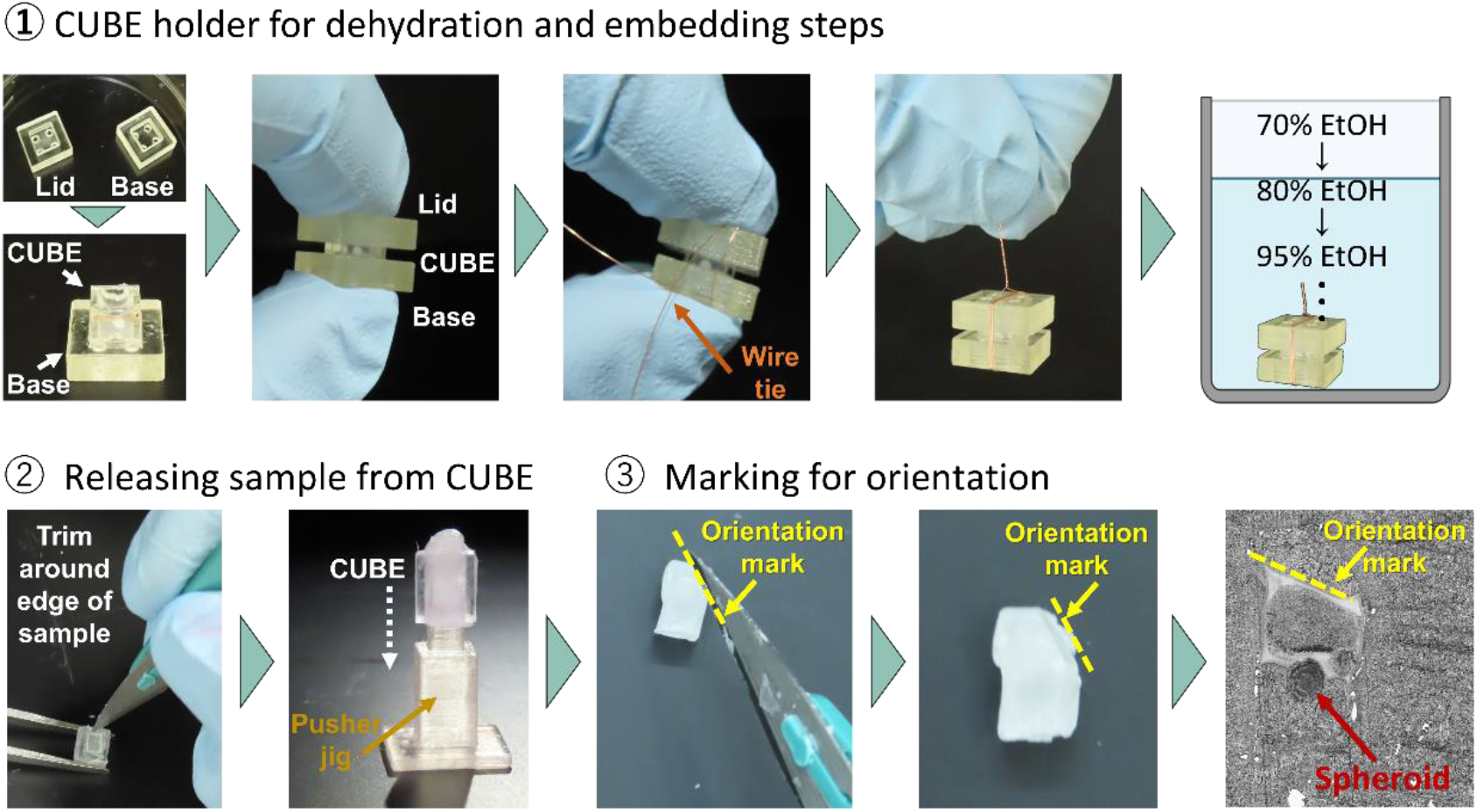
Detailed process showing the preparation of sample for paraffin sectioning. A CUBE holder was designed to prevent the risk of losing samples during the paraffinization process. After paraffinization, the sample is released from the CUBE and marked for orientation by cutting one edge of the sample.

**Supplementary Figure 3.**
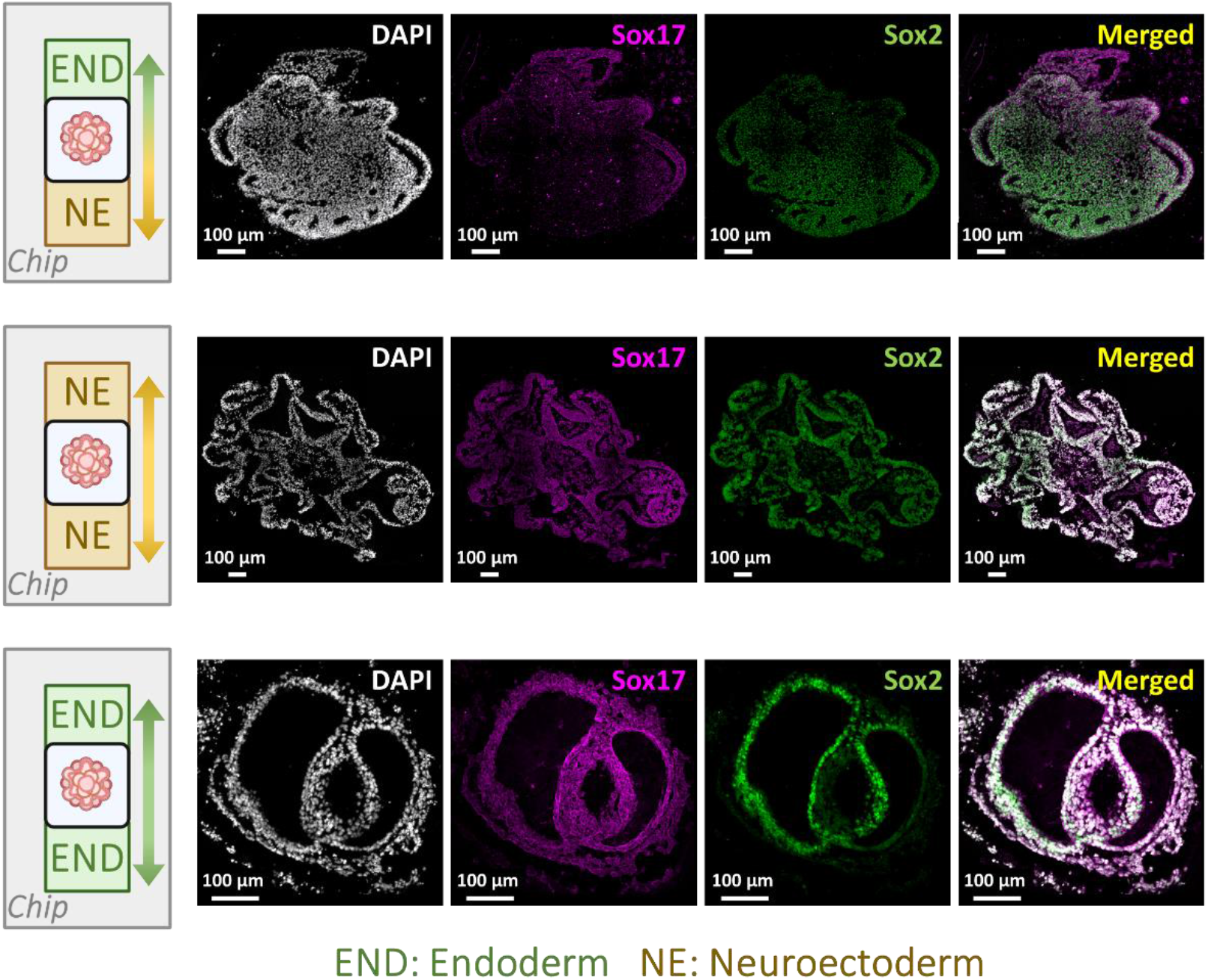
Immunofluorescence of paraffin-sectioned sample of END-NE differentiated spheroid. Immunostaining showed localized expression pattern of endoderm (Sox17) and neuroectoderm (Sox2) markers in END-NE spheroids, whereas NE only and END only controls showed uniform distribution of the markers.

